# Artificial Intelligence Apps for Medical Image Analysis using pyCERR and Cancer Genomics Cloud

**DOI:** 10.1101/2025.01.19.633756

**Authors:** Aditya P. Apte, Eve LoCastro, Aditi Iyer, Sharif Elguindi, Jue Jiang, Jung Hun Oh, Harini Veeraraghavan, Amita Shukla-Dave, Joseph O. Deasy

**Affiliations:** Department of Medical Physics, Memorial Sloan Kettering Cancer Center, New York, USA

## Abstract

This work introduces a user-friendly, cloud-based software framework for conducting Artificial Intelligence (AI) analyses of medical images. The framework allows users to deploy AI-based workflows by customizing software and hardware dependencies. The components of our software framework include the Python-native Computational Environment for Radiological Research (pyCERR) platform for radiological image processing, Cancer Genomics Cloud (CGC) for accessing hardware resources and user management utilities for accessing images from data repositories and installing AI models and their dependencies. GNU-GPL copyright pyCERR was ported to Python from MATLAB-based CERR to enable researchers to organize, access, and transform metadata from high dimensional, multi-modal datasets to build cloud-compatible workflows for AI modeling in radiation therapy and medical image analysis. pyCERR provides an extensible data structure to accommodate metadata from commonly used medical imaging file formats and a viewer to allow for multi-modal visualization. Analysis modules are provided to facilitate cloud-compatible AI-based workflows for image segmentation, radiomics, DCE MRI analysis, radiotherapy dose-volume histogram-based features, and normal tissue complication and tumor control models for radiotherapy. Image processing utilities are provided to help train and infer convolutional neural network-based models for image segmentation, registration and transformation. The framework allows for round-trip analysis of imaging data, enabling users to apply AI models to their images on CGC and retrieve and review results on their local machine without requiring local installation of specialized software or GPU hardware. The deployed AI models can be accessed using APIs provided by CGC, enabling their use in a variety of programming languages. In summary, the presented framework facilitates end-to-end radiological image analysis and reproducible research, including pulling data from sources, training or inferring from an AI model, utilities for data management, visualization, and simplified access to image metadata.

## 1 Introduction

This work addresses the need for a comprehensive cloud-based software framework for medical image analysis. Data and AI models are persistently hosted in linked cloud buckets from providers such as AWS and GCP or pulled in from external cloud repositories, including public sources such as TCIA^3^ and IDC^4^ as well as proprietary sources such as XNAT^7^, BOX (https://www.box.com). Cancer Genomics Cloud (CGC)^8, 9^ provides integrated bioinformatics tools, automatic logging, versioning, and access-controlled sharing, facilitating reproducibility and collaboration. Figure 1 shows various components of the cloud-based framework for end-to-end AI workflows for medical image analysis.

**Figure 1:**
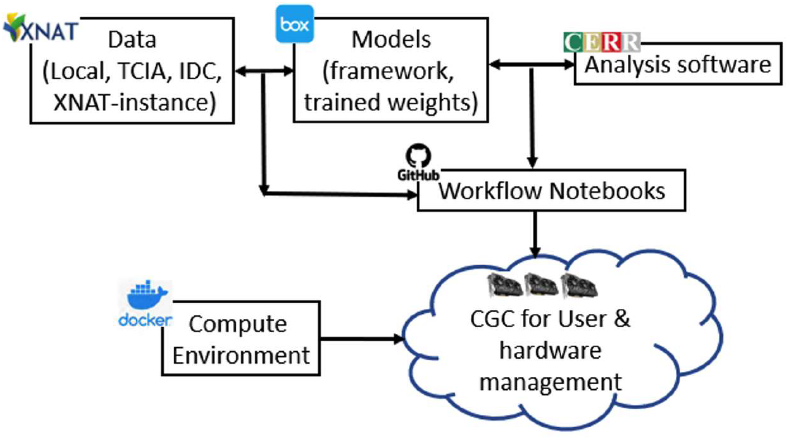
Components of cloud-based analyses for medical imges

**Figure 2:**
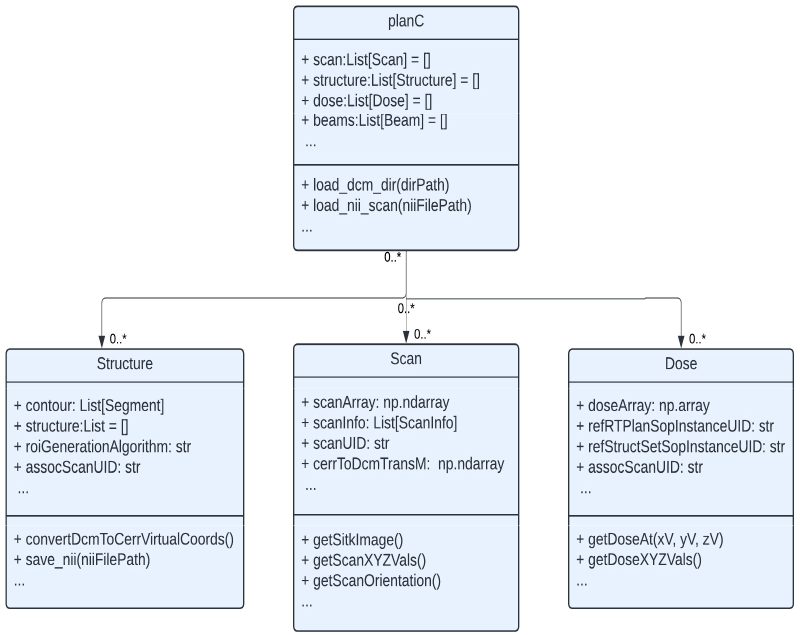
pyCERR data classes. The planC container class is extensible and composed of different radiological data types.

The Computational Environment for Radiological Research (CERR) platform^13^, developed in the MATLAB programming language has been widely used and adapted for various medical physics applications over the past decade due to its robust and user-friendly data structure and visualization capabilities. These include prototyping Intensity Modulated Radiation Therapy (IMRT) optimization algorithms^14^, outcomes modeling^15, 16^, film dosimetry QA^17^, quality assurance of RT^18^ and imaging data by leading institutions, and CNN-based training and inference^15^. CERR’s flexible data structure has been extended by several plugins such as segmentation benchmarking software developed by AAPM TG 211^16^. While CERR has been highly useful in clinical research, its adoption is limited by the requirement of MATLAB license for development. This is especially crucial in cloud-environments where open-source programming languages offer license-free use and readability of the underlying codebase. Python has emerged as the language of choice for data analysis and research, serving as a glue language for various software libraries. Python’s scientific computing and visualization libraries now offer functionalities that are on par or surpass MATLAB and its toolboxes for medical image processing.

pyCERR maintains the flexible, readable data structure and graphical user interfaces of the CERR platform while providing relevant tools in the Python programming environment. This allows pyCERR to seamlessly integrate with the medical image analysis ecosystem and software such as XNAT, 3DSlicer, ANTPy, etc. enabling access to cutting-edge AI modeling and image processing libraries for local and cloud-friendly workflows. pyCERR is used in clinical deployment of AI segmentation models at our institution which have been made publicly available for research use.

We describe various AI-based apps developed using pyCERR and deployed on the CGC. This framework fosters collaborative research through persistent hosting of data, code, and analysis tools on the cloud. It enhances the medical image analysis ecosystem by using leveraging data archival platforms, libraries to train and infer AI models, cloud providers and Python APIs from popular image analysis libraries^19, 20^. The next sections describe the architecture and various modules of our cloud-based framework in detail.

## 2 Materials and Methods

### (i) Data I/O

Data analysis workflows require radiological inputs (images, doses, segmentations) and AI models. Radiological images are hosted in publicly available repositories such as TCIA, IDC, Zenodo (https://zenodo.org) which provide Python APIs to query/retrieve images. We have hosted an instance of XNAT (“PIXNAT”) at our institution which is accessible to external collaborators to archive images for research. <u>It</u> facilitates the sharing of IRB-approved project datasets with external collaborators. XNAT provides APIs (pyXNAT, XNATpy) to query, retrieve, and upload images.

Trained AI models consist of the network architecture and associated weights. Depending on the network architecture, the weight files can be substantial in size. We archive the trained network along with its dependencies in the form of a packaged Anaconda environment. The network and weights are stored in our institution’s BOX platform with publicly available static links for direct download.

Additionally, cloud providers such as GCP and AWS provide data storage which can be accessed programmatically. Users are charged for hosting as well as egress to move data to and from these buckets.

### (ii) Analysis software

The components of pyCERR can be broadly categorized as (1) Data container, (2) Viewer, and (3) Analysis modules. pyCERR builds on commonly used scientific libraries Numpy, scipy, pandas and image processing libraries such as pydicom, SimpleITK, and scikit-image. pyCERR uses Napari for visualization via graphical user interface objects within the Python environment.

#### (a) Metadata container

The central idea of pyCERR is to provide a consistent data structure for metadata from different file formats. The plan container class “planC” houses data classes containing attributes for various data types. These include (a) Scan for imaging data such as CT, MR, PET, (b) Structure for image segmentation, (c) Dose for RT dose, and (d) Beams for RT plans as well as derived data types such as for deformable registration. Figure 1 shows a schematic representation of data classes, including key attributes and routines for data access and transformation.

Modules are provided for importing and exporting imaging data between DICOM and NifTI formats as well as conversion to SimpleITK^21^ Image objects. Pydicom^22^ is used to read and write DICOM, Nibabel^23^ to write to NifTI, and SimpleITK to read ITK-compatible formats such as NifTI, MHA, and NRRD. Data attributes in the planC container can be readily accessed for analyses, edited, and saved to DICOM and NifTI file formats.

pyCERR currently imports metadata from DICOM RTSTRUCT and SEG segmentation modalities, scan modalities CT, MR, PET and radiotherapy dose modalities RTDOSE and RTPLAN. Segmentations are stored as polygonal contours and can be converted to binary masks and vice-versa. pyCERR provides utilities to convert PET images to SUV^24^ units using normalizations such as Body Weight, Body Surface Area, and Lean Body Mass approaches. pyCERR supports dynamic scan sequences such as DCE and diffusion weighted MRI, with image volumes are organized by acquisition time or b-value. MRI images are converted to real world units using vendor-specific private tags.

#### (b) Visualization

Multi-dimensional image viewer Napari^25^ is perfectly suited for pyCERR’s multi-modal data structure. As Napari is written in, it eliminates the need for users to transfer data between viewers written in other languages, such as ImageJ^26^ and 3D Slicer^27^. The Napari ecosystem provides a rapidly-growing list of plugins for image analysis, such as formicroscopy datasets and the Segment Anything Model^28^. To our knowledge, pyCERR is the only software platform that enables visualization of radiotherapy data in Napari. The viewer module in pyCERR provides routines to visualize scan, segmentation, and dose data types. It provides widgets to window scans and to segment them by leveraging Napari’s labeling tools.

#### (c) Analysis modules

Organization of imaging data in a plan container enables efficient access to the underlying data elements and their relationships for building analysis modules. Modules for data transformation, radiomics, dosimetric analysis, and segmentation evaluation are included with pyCERR. Data transformations include tools for post-processing segmentations, such as smoothing contour polygons, connected components filtering, resampling and transferring segmentations and doses from one scan grid to another. The radiomics module in pyCERR provides scalar features and texture filters consistent with their IBSI definitions. Feature and filter computations are entirely Python-based, using vectorized code via Numpy arrays for speed. Implementations were validated against IBSI benchmarks. IBSI2 Filter response maps are stored in planC, which facilitates visualization and analysis. pyCERR provides tools to extract dose-volume histogram (DVH)-based features such as Vx, Dx, MOHx, MOCx.

### (iii) User management and sharing

User data and workflows are organized into projects on CGC; they can be kept private or made available for sharing with other users on the platform. Users create and share analysis workflows as “apps” that are launched in user-specified Docker environments. Apps are defined in Common Workflow Language (CWL); multiple CWL tasks can be chained together in a workflow definition. These workflows can be initiated as tasks in the cloud within the CGC-SB web portal or externally from users’ local hardware using their API via Python or command line.

CGC-SB Interactive Data Studio environments include Jupyter, RStudio, SAS Studio and Galaxy. The OHIF^29^ viewer is also available for reviewing images in the project. CGC Workflow apps and Data Studio operate on AWS instances with specific CPU, RAM or GPU requirements configured by user.

## 3 Results

AI workflows are organized as a sharable project on CGC platform. Pretrained AI models are provided for segmentation of normal tissues across multiple sites. Table 1 lists models available on the platform. These can be accessed via CGC Seven Bridges API or via an interactive Jupyter notebook. A locally runnable Jupyter notebook demonstrating API call to heart sub-structure segmentation model is distributed at https://github.com/cerr/pyCERR-Notebooks/blob/main/SBG_autosegment_CT_Heart_OARs.ipynb.

**Table 1:**
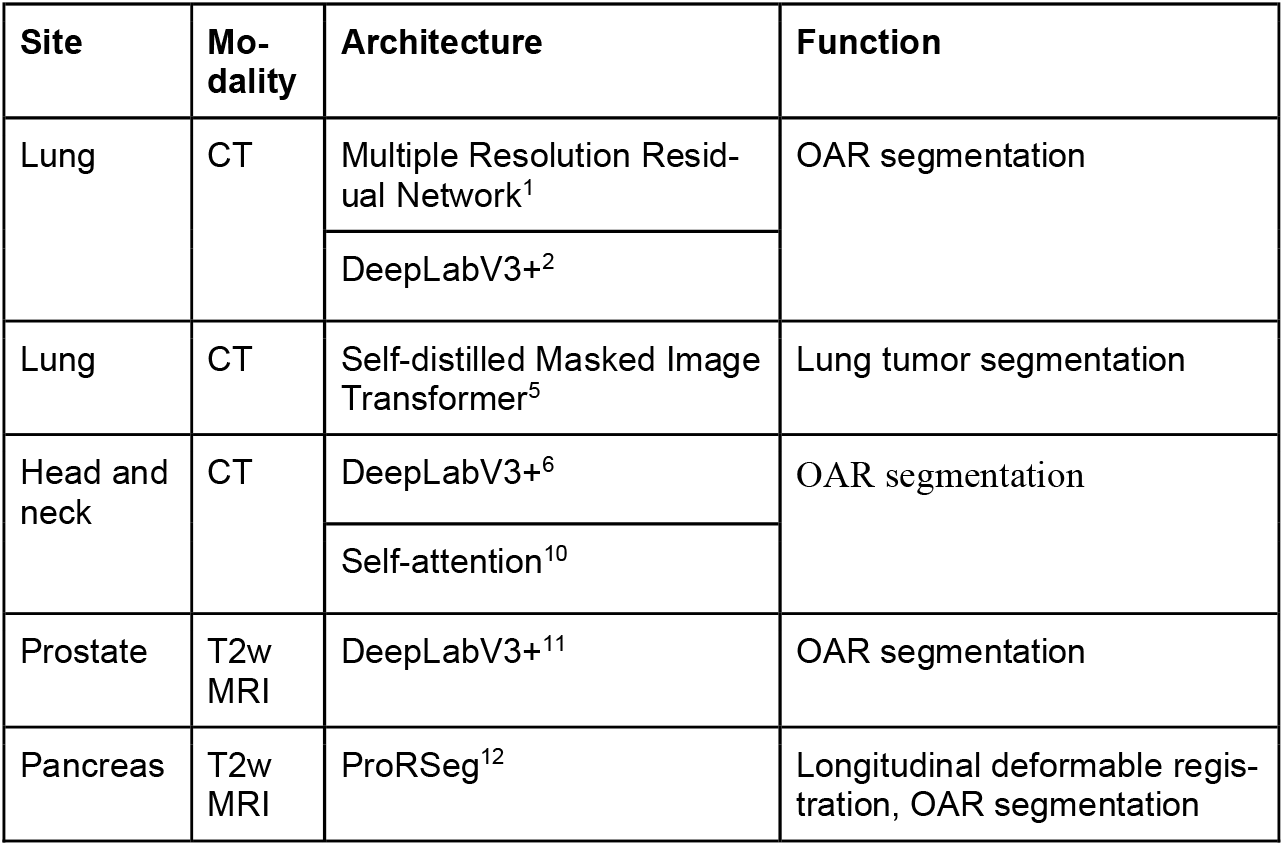
Pre-trained AI models

The pyCERR package is distributed under an open-source, GNU GPL license at https://github.com/cerr/pyCERR. pyCERR can be installed from the GitHub repository by using Python package installer pip. Tests for various modules including IBSI-compliant radiomics, and conversion of data between various formats are distributed with pyCERR. These are run upon every code update to the GitHub repository. Notebooks demonstrating various use cases are provided at https://github.com/cerr/pyCERR-Notebooks to provide a starting point for analyses. Following are some common use cases:

### (1) Re-Radiation / Adaptive RT, dose accumulation

pyCERR’s data structure supports multi-modal and longitudinal imaging data, facilitating dose accumulation for re-radiation and adaptive RT by transferring segmentations and dose distributions between different time-points via image registration. Visualization tools are provided for quality assurance of image registration as well as deformation vector field (DVF). Figure 3 shows an example of mirrorscope and DVF display.

**Figure 3:**
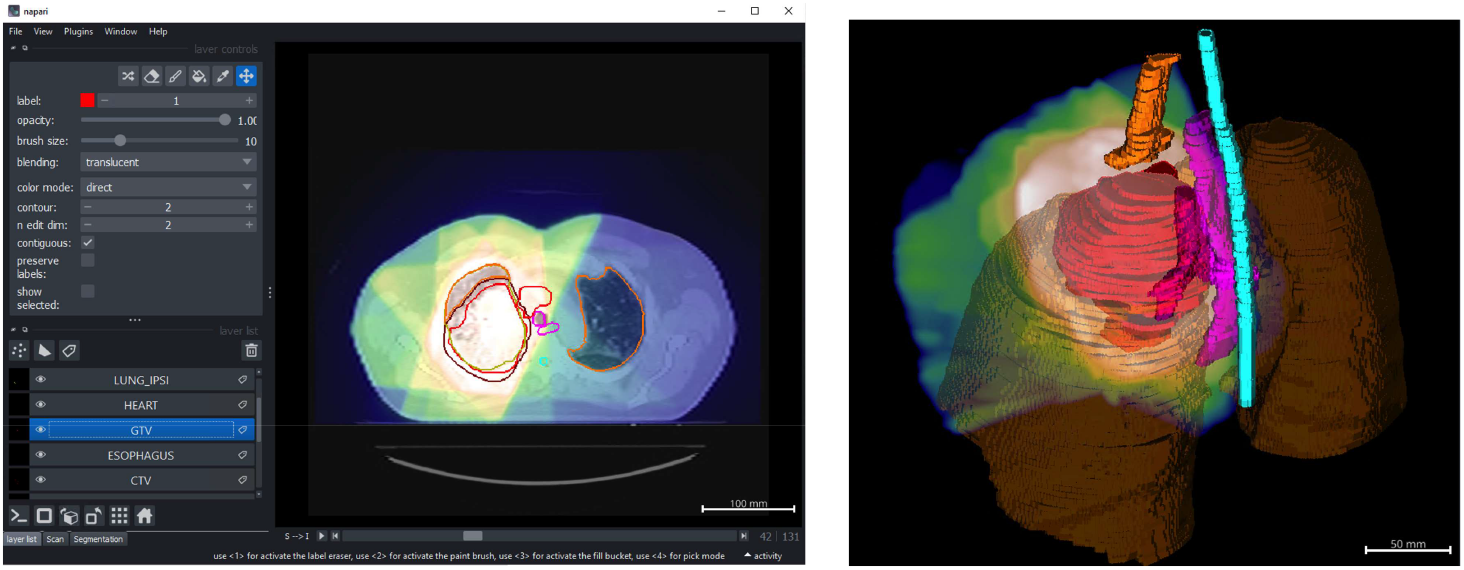
Radiotherapy does distributions can be visualized along with structures in 2D and 3D. Display settings such as Window, transparency, gamma can be adjusted from GUI or command, thus making it suitable for capturing screenshots in batch mode for large number of datasets.

### (2) Visualizing a batch of datasets

pyCERR’s viewer module provides routines to overlay imaging data from the planC container such as scans, doses and structures as demonstrated in figure 3. These routines can be used locally in an interactive mode or on a remote server in batch-mode; The notebook at https://github.com/cerr/pyCERR-Notebooks/blob/main/batch_visualize_scan_seg_ex1.ipynb demonstrates capturing screenshots for a batch of DICOM datasets on Google Colab platform.

### (3) Image segmentation

The cloud-based framework provides well-validated tools for normal tissue segmentation for various radiotherapy treatment sites as listed in Table 1. Additionally, pyCERR provides a manual segmentation widget supports multiple orthogonal views, overlaying image modalities such as PET and CT, as demonstrated in Figure 4. This can be used to generate ground-truth data for model training as well as editing the output of AI segmentation for downstream use.

**Figure 4:**
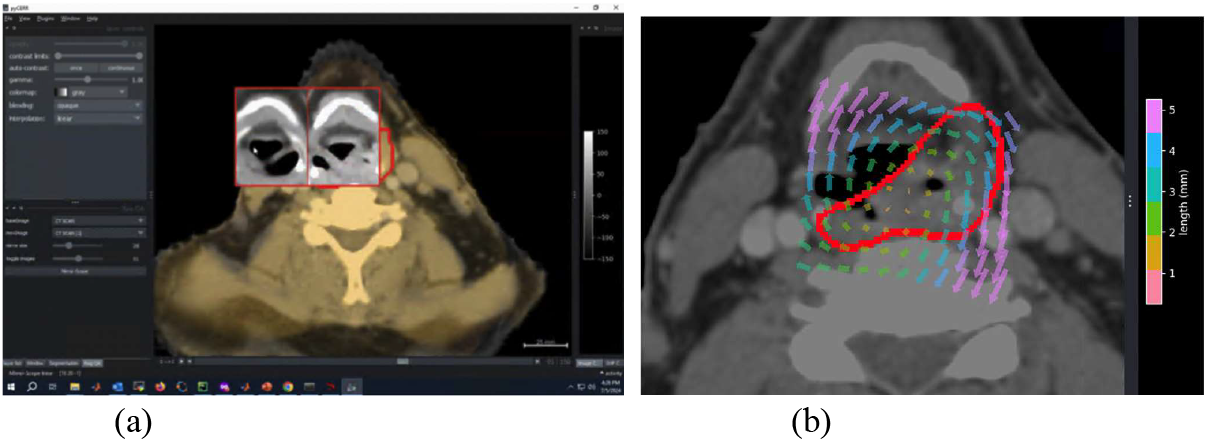
Mirrorscope and DVH display

**Figure 5:**
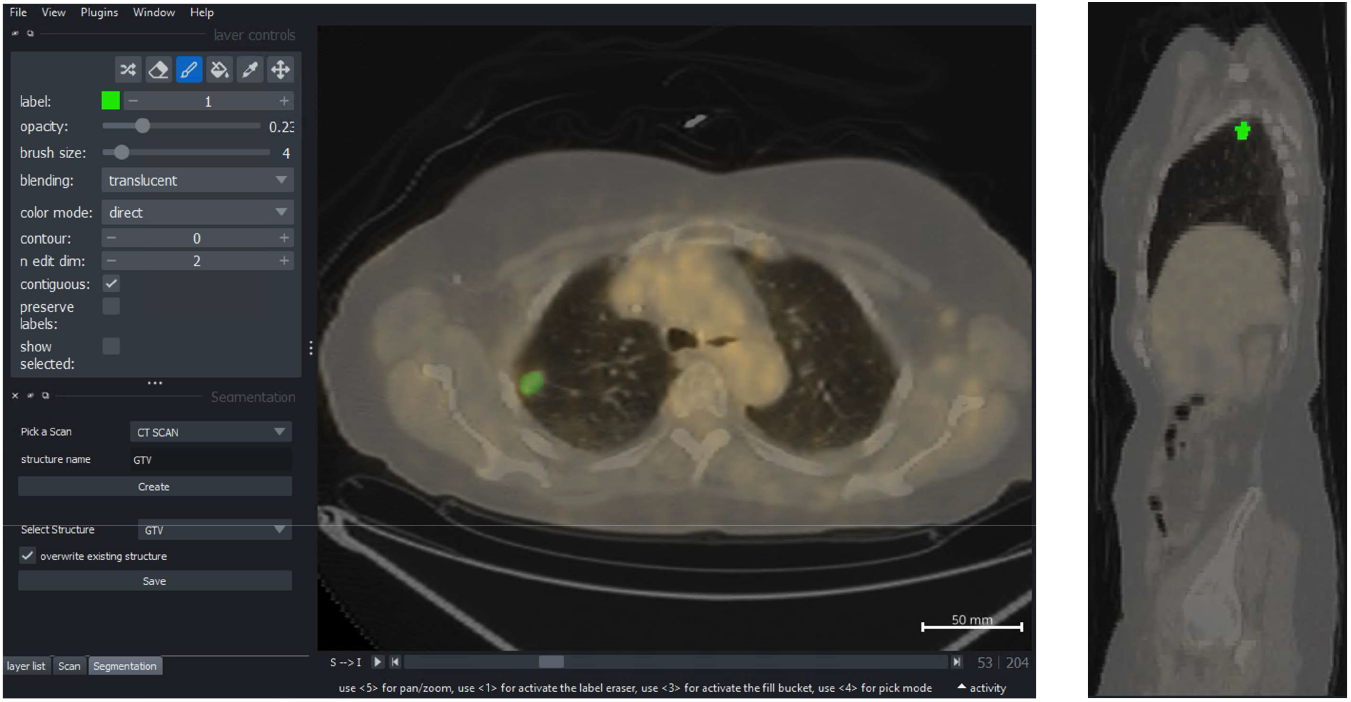
Contouring tools in pyCERR. The segmentation widget provides the ability to create and edit structures. This is useful to edit contours resulting from auto-segmentation as well as to generate new structures. Napari provides Paintbrush, Eraser and Fill tools for editing segmentation labels.

**Figure 6:**
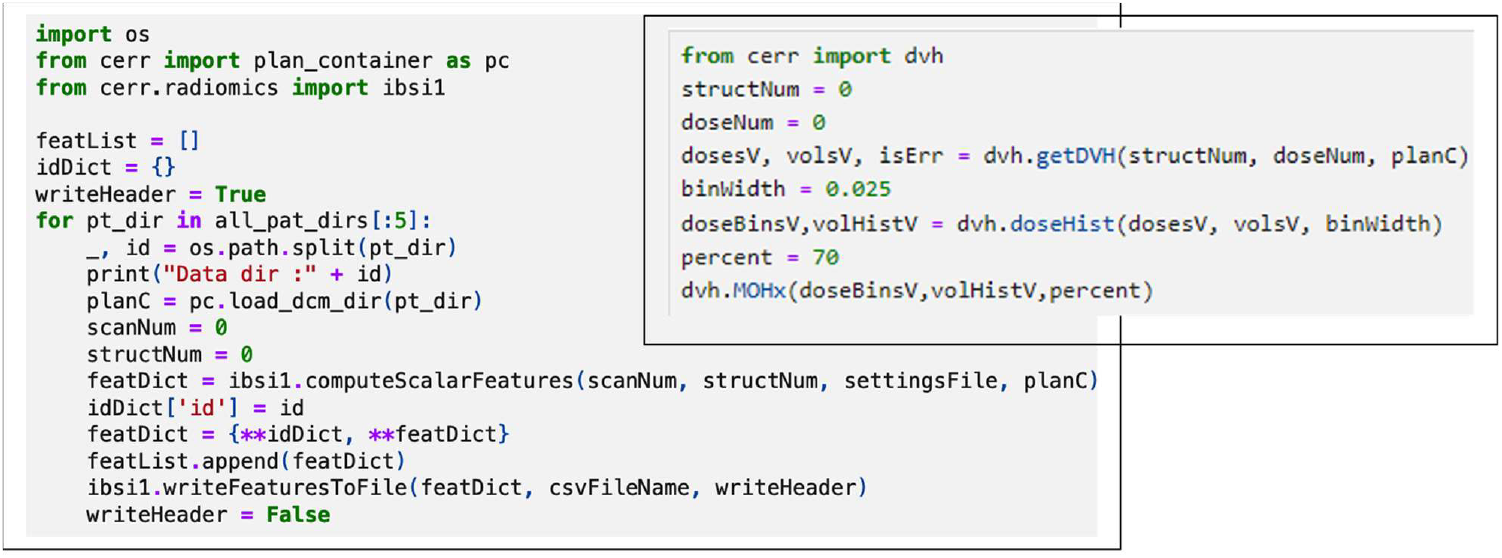
(a) Radiomics feature extraction for a batch of datasets: The ***radiomics*** module provides the flexibility of selecting scan and segmentation objects from planC and specifying calculation settings via JSON (b) DVH feature extraction: The ***dvh*** module provides routines to obtain RT dose within the segmentation, accumulation of dose values into histograms and calculation of scalar metrics.

### (4) Radiomics and DVH-based metrics calculation

pyCERR provides “*radiomics*” and “*dvh*” modules to support feature extraction from scan and dose datasets. The *radiomics* module provides routines for IBSI-1 and IBSI-2 compatible feature extraction. The dvh module provides an ability to compute commonly used metrics with user defined dose bin width.

## 4 Discussion

The presented framework simplifies the use of AI models. Data processing is pre-configured using pyCERR, and hardware dependencies are easily setup on the CGC, extending access to users with minimal programming experience. Moreover, users can readily combine imaging with genomics data available on CGC.

Napari is a relatively new project compared to more established viewers such as ITK-SNAP^30^ and 3DSlicer. While the underlying Napari features are undergoing continuous development, it is the only Python-based viewer with a rich feature set. Napari, as a generic image viewer, supports visualization of various data types such as shapes, vectors, labels, images and allows for defining custom callback events. This enables users to customize pyCERR’s visualization tools to suit their specific needs. Additionally, pyCERR will benefit from future development in Napari, including support for orthogonal view planes, grouping of display layers, and viewing remote datasets.

## 5 Conclusion

The framework for deploying AI apps presented in this work is accomplished without local setup or installation required in basic command line tools to remotely launch apps on the CGC platform. pyCERR provides comprehensive tools for radiological and radiotherapy data processing, leveraging open-source formats and libraries. Users are provided with a shareable project on CGC to quickly get started on using the presented analyses. The tools and Jupyter notebooks are self-contained such that they can also be run on institutional HPCs or other cloud platforms.

## Acknowledgements

This study was supported in part by the MSK Cancer Center Support grant (NIH grant P30 CA008748), The Simons Foundation, the Breast Cancer Research Foundation (grant MATH-23-001), and the NIH ROBIN cooperative group (grant U54CA274291). We also acknowledge in-kind support for cloud computation from Seven Bridges Cancer Research Data Commons Cloud.

The Seven Bridges Cancer Research Data Commons Cloud Resource has been funded in whole or in part with Federal funds from the National Cancer Institute, National Institutes of Health, Contract No. HHSN261201400008C and ID/IQ Agreement No. 17×146 under Contract No. HHSN261201500003I and 75N91019D00024.

